# Allosteric coupling between α-rings of the 20S proteasome

**DOI:** 10.1101/832113

**Authors:** Zanlin Yu, Yadong Yu, Feng Wang, Alexander G. Myasnikov, Philip Coffino, Yifan Cheng

**Affiliations:** Department of Biochemistry and Biophysics, University of California San Francisco, CA, 94158; Laboratory of Cellular Biophysics, Rockefeller University, New York, NY, 10065; Howard Hughes Medical Institute, University of California San Francisco, San Francisco, CA 94158

## Abstract

The proteasomal machinery performs essential regulated protein degradation in eukaryotes. Classic proteasomes are symmetric, with a regulatory ATPase docked at each end of the cylindrical 20S. Asymmetric complexes are also present in cells, either with a single ATPase or with an ATPase and non-ATPase at two opposite ends. The mechanism that populates these different proteasomal complexes is unknown. Using archaea homologs, we constructed asymmetric forms of proteasomes. We demonstrate that the gate conformation of two opposite ends of 20S are coupled: binding one ATPase opens a gate locally, and also the remote opposite gate allosterically. Such allosteric coupling leads to cooperative binding of proteasomal ATPases to 20S, and promotes formation of proteasomes symmetrically configured with two identical ATPases. It may also promote formation of asymmetric complexes with an ATPase and a non-ATPase at opposite ends. We propose that in eukaryotes a similar mechanism regulates the composition of the proteasomal population.

## Introduction

In eukaryotic cells, 26S proteasomes are complex protein machines that execute regulated and ATP-dependent intracellular protein degradation. Its functions include degrading and thus detoxifying misfolded proteins and controlling the amounts of regulatory proteins, such as those involved in cell cycle regulation ^1^. The 26S proteasome is composed of the 20S core particle (CP), symmetrically capped at each end by a 19S regulatory particle (RP). The proteolytic module, the 20S CP, is a hollow barrel-shaped chamber formed by four stacked heptameric rings, two inner β-rings and two outer α-rings. The proteolytic active sites at N-termini of the β-subunits are sequestered within the chamber ^2,3^. Unfolded substrates enter the chamber through a narrow pore in the center of α-rings, where N-termini of α-subunits form a gate to impede unregulated access to the proteolytic chamber ^4^. The primary role of 19S RP is to recognize, unfold and translocate targeted substrates into CP for degradation ^5^. In addition, the six proteasomal ATPases (Rpt subunits) within 19S RP also engage with the α-ring of 20S CP and open its gate for unfolded substrates to enter the proteolytic chamber and, presumably, for small peptide products of proteolysis to be released from the chamber ^6^.

The canonical 26S proteasome has a 2-fold symmetrical architecture, with a 19S RP cap at each end of the 20S CP. CPs capped by a single 19S RP, or uncapped, are also abundant in cells ^7^. Other types of nonATPase RP complexes are also present in eukaryotes, such as PA28 and PA200 ^8,9^. Thus, there are hybrid particles, in which 20S CP is capped at each end by a different RP ^7,10-14^. The functions of these non-ATPase activators in the regulated protein degradation process remain elusive, but invariably include opening the α-ring gate. Consequently, as defined by alternative cap configurations, proteasomes constitute a diverse family. A key question is in what way these structural alternatives differ in their biochemical capacities and cellular functions. More specifically, how do 20S CP symmetrically capped with two 19S RP compare with those bearing a single 19S RP? Is one cap sufficient to make the proteasomal particle fully functional? Furthermore, are the effects of RP docking on the α-ring purely local, or do such effects propagate distantly? Such questions are not readily addressed with eukaryotic 26S proteasomes, because it is difficult to biochemically separate canonical symmetric 26S particles from those with only one 19S RP for functional studies.

In this study, we use archaeal proteasomes to examine these long-standing questions. Specifically, we ask whether substrate processing efficiency is enhanced by the presence of two RPs and whether gate opening can be rate determining. As a model system, the archaeal proteasome is similar to but simpler than eukaryotic proteasomes ^15^. In archaea, the 20S CP is composed of homo-heptameric α- and β-rings ^3^, and PAN (Proteasomal Activator Nucleotidase) is a homohexamer proteasomal ATPase found in many species ^16,17^. Archaeal PAN ATPase, like the ATPase ring of eukaryotic 19S CP, docks to the α ring via a C-terminal HbYX (hydrophobic, tyrosine, any amino acid) motif ^18^. In the past, many mechanistic insights relevant to eukaryotic 26S proteasome were derived from studies of archaea proteasome ^16-19^.

We designed and assembled *in vitro* various asymmetric archaea 20S CP, in which the two individual α-rings contain different point mutations, so that only one α-ring can bind to a specific RP. This approach enabled us to measure and compare proteolytic activities of singly capped 20S CP versus doubly capped ones, and to reveal structurally evident allosteric coupling between the two separate α-rings upon RP binding. Our studies reveal that, upon binding of a RP with a conserved C-terminal HbYX motif to one α-ring, the gate of the opposite α-ring opens allosterically, resulting in similar proteolytic activities of 20S CP singly capped and doubly capped with PAN. Such allosteric regulation of the α-ring gate requires the C-terminal HbYX motif be present in RP. Furthermore, allosteric coupling between two α-rings leads to cooperative binding of activator to 20S CP. We hypothesize that there is a similar allosteric coupling between the two α-rings in eukaryotic 26S proteasomes.

## Results

### *In vitro* assembling of asymmetrical archaeal 20S proteasome

Functional archaeal 20S CPs can be assembled *in vitro*^20^ by mixing and incubating purified full length α- and pro-peptide truncated β-subunits that are individually expressed in *E. coli* (Supplementary Figure 1). When expressed and purified alone, α-subunits assemble into homo-heptameric rings. Two α-rings come together back-to-back, forming a tetradecameric double α-ring, with the α-subunits in each ring interdigitated with other α-subunits, similar to the manner in which they interdigitate with β-subunits in an intact native 20S CP (Supplementary Figure 1C-I and Supplementary Table 1). Mixing such double α-rings with β-subunits, which remain as monomers throughout the purification, induces separation of the α tetradecamer double rings into two heptameric α-rings and results in the assembly of α- and β-subunits together into intact 20S CPs. Negative stain electron microscopy (EM) shows *in vitr*o assembled 20S CPs are indistinguishable from the native recombinant 20S CP assembled *in vivo* (Supplementary Figure 1J). We also confirmed that the *in vitro* assembled 20S CPs are functional in terms of proteolytic activity and activation of α-ring gating by proteasomal activator, although its proteolytic activity level is slightly lower than that of the native 20S CP (Supplementary Figure 1K).

Because α-rings can be pre-assembled homogeneously with the same type of α-subunit, it is possible to mix two different types of pre-assembled α-rings with β-subunits to assemble 20S CP with two different types of α-ring at its opposite ends. We generated two different types of α-rings with different affinity tags: one was made by co-expressing two full length α subunits in one expression plasmid, one with a His-tag and the other with an MBP-tag, both at the C-terminus. Such assembled α-rings thus contain a mixture of two different affinity tags to facilitate multi-step affinity purification. A second type of α-ring was made with a single Strep-Tactin tag. (We found that Strep-Tactin tag can be used multiple times for affinity purification, but His-or MBP-tags can only be used once.) The β-subunit uniformly had a C-terminal His-tag. A TEV cleavage site was placed before all affinity tags. We first purified pre-assembled His/MBP-tagged α-rings by a Ni-NTA affinity column, and the Strep-Tactin-tagged α-rings by a strep-trap affinity column. β-subunits were purified by using a Ni-NTA affinity column. We then assembled intact CP proteasomes *in vitro* by mixing the pre-assembled α-rings and β-subunits. The resultant CP population is a mixed population, some portion of which constitute the desired asymmetric CPs with distinct α-ring at each end (Figure 1A). To purify these, a two-step affinity purification scheme was used, by Strep-trap tag first, followed by MBP tag. By this means, purified 20S CPs that contain two different types of α-rings were obtained (Figure 1A). After purification, all affinity tags were removed by TEV protease cleavage.

**Figure 1.**
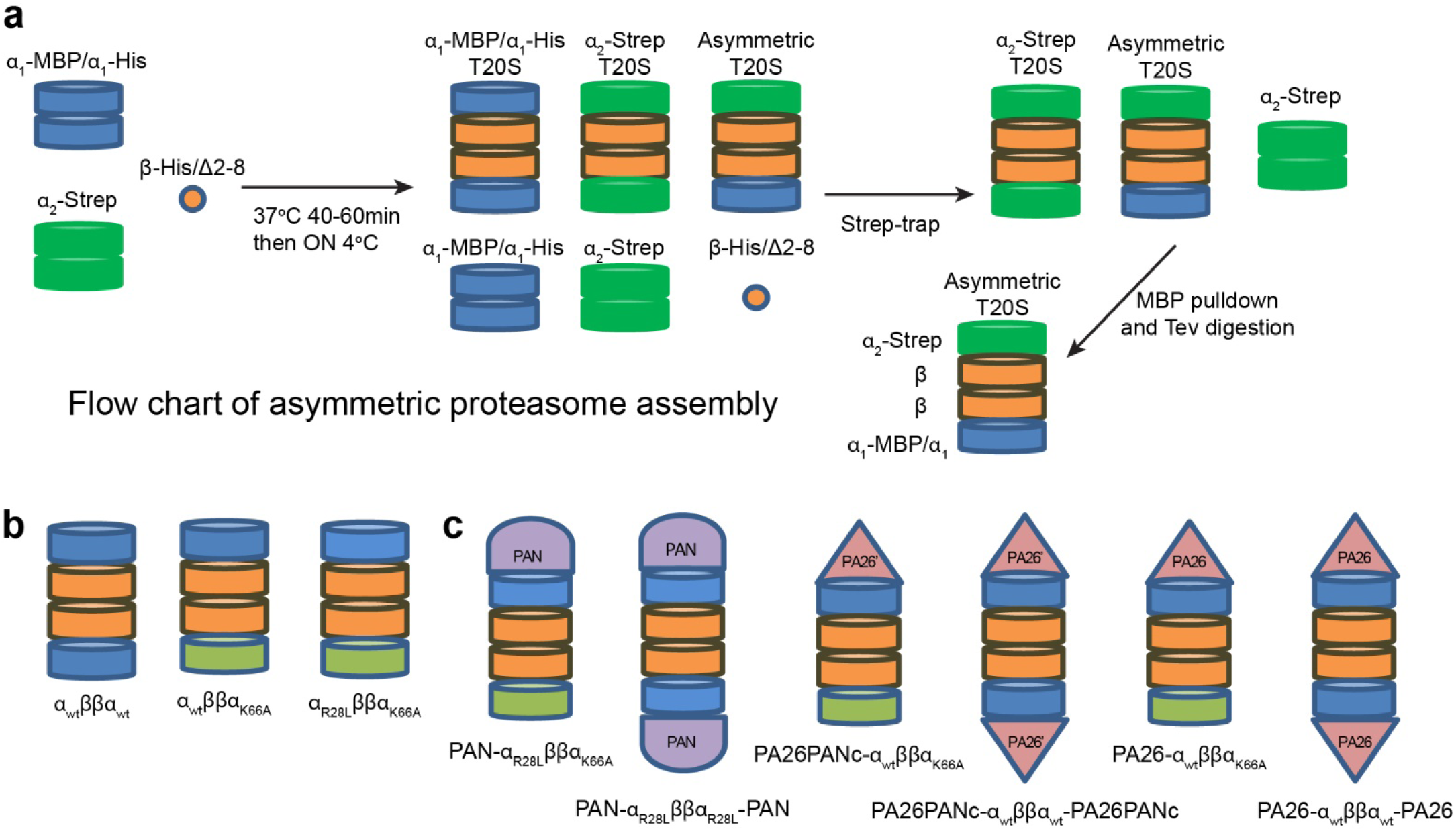
Scheme for in vitro assembly of asymmetric proteasome. **A**: Schematic flowchart of asymmetric 20S CP *in vitro* assembly and purification. Following assembly of the depicted mixed population, the intended asymmetric particles were purified by a two-step affinity purification scheme. **B**: Schematic illustration of the types of *in vitro* assembled asymmetric 20S CPs used in this study. **C**: Schematic illustration of all types of RP-capped complexes assembled and used in this study.

Using this strategy, we assembled 20S CPs that contain two distinct α-rings at their opposite ends. One end is wild type and the other contains the point mutation K66A. Such asymmetric 20S CP, named α_WT_ββα_K66A_, allow the binding of an activator to the wild-type α-ring but not to the opposite α-ring with the K66A mutation, which impairs binding of any activators including both proteasomal ATPase with a C-terminal HbYX motif or 11S activators such as PA26 from *Trypanosoma brucei*^18,21^. This allowed formation of a homogeneous population of singly capped proteasomes. As a control, we also assembled *in vitro* symmetric wild type 20S CP α_WT_ββα_WT_ for comparison. A full listing of *in vitro* assembled complexes is shown in Figure 1B.

In this study, we used three types of activators. The first one is a wild type homo-hexameric PAN ATPase, homologous to the Rpt ATPase subunits in the base of 19S RP of eukaryotic 26S proteasomes. The C-terminus of PAN contains an HbYX (hydrophobic, tyrosine, any amino acid) motif that is required for binding to the α-ring and opening its gate ^18^. When PAN is used, we also introduced a point mutation R28L to the corresponding α-ring of 20S CP. This mutation does not influence the gating or proteolytic activity but enhances binding of PAN to CPs. The second activator is heptameric PA26 with a point mutation of E102A and the last 8 C-terminal residues replaced with the last 8 C-terminal residues from PAN (with HbYX motif), which we named PA26PANc. PA26 is a heptameric 11S activator from *Trypanosoma brucei*^22^. The point mutation E102A abolishes a specific interaction between the so-called activation loop of PA26 with a reverse turn loop in the 20S α-subunit, thus abolishing the capability of PA26 to activate 20S proteasomes by opening its gate. Instead, the HbYX motif from the PAN C-terminus placed at the heptameric PA26PANc C-termini performs gate opening ^23^. As demonstrated previously, PA26PANc activates the CP particle in the same manner as PAN, except that it symmetrically engages the seven-fold symmetric α-ring, and thus avoids a symmetry mismatch between the α-ring and the activator ^19,23^. The third activator is PA26 with a single point mutation of V230F, which functions like the wild type PA26, but with an enhanced binding affinity to the archaeal 20S CP ^23^. Mixing our *in vitro* assembled CP with these activators generated the desired types of single or double capped proteasomes (Figure 1C and Supplementary Figure 2).

**Figure 2.**
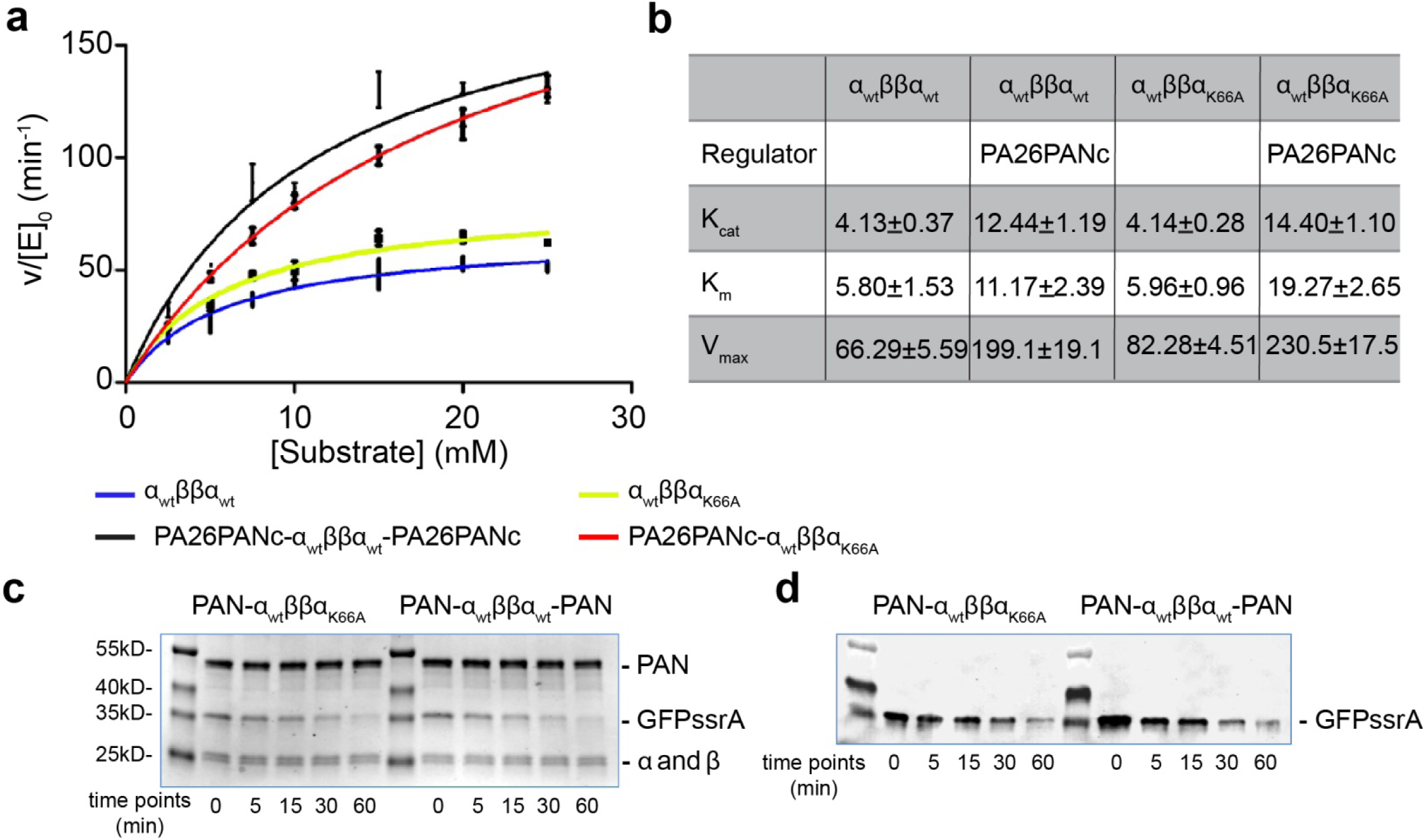
Substrate degradations by asymmetric proteasomes. **A**: Kinetics of LFP peptide degradation by symmetric (blue) and asymmetric (yellow) 20S CP alone and with double (black) and single (red) activator PA26PANc. **B**: Table summarizing the *k*_*cat*_, *K*_*M*_, and *V*_max_ of LFP peptide degradation by symmetric and asymmetric 20S CP without and with activators. **C**: Time course of GFP-ssrA degradation by PAN-α_WT_ββα_K66A_ and PAN-α_WT_ββα_WT_-PAN, analyzed by Coomassie Blue stained SDS PAGE. **D**: Time course of GFP-ssrA degradation by PAN-α_WT_ββα_K66A_ and PAN-α_WT_ββα_WT_-PAN, analyzed by Western blotting with anti-GFP antibody.

### Proteolytic activities of asymmetric 20S proteasome

We aimed to use the *in vitro* assembled asymmetric 20S CP to address the question of whether the singly capped proteasome has the full peptidase or protease capacity of a doubly capped one. To this end, we used two types of substrate, LFP and GFP-ssrA. LFP is a fluorogenic peptide of nine amino acids that emits florescence upon proteolytic cleavage. It can diffuse into the archaeal 20S proteasome and undergo proteolysis even without an activator, but gate opening increases its entry rate. Upon opening the α-ring gate, fluorescent product is seen to accumulate at a faster rate, consistent with an increased flux of LFP peptides into the 20S degradation chamber ^18^. GFP-ssrA is a folded protein substrate containing the very stable and bulky GFP β-barrel, which cannot passively diffuse into the 20S CP proteasome for degradation, even when its gate is open. Degradation of GFP-ssrA requires a 20S CP in complex with PAN ATPase; PAN hydrolyzes ATP to unfold GFP-ssrA and to actively translocate the unfolding polypeptide into the CP, where proteolysis takes place.

Using PA26PANc as the activator, we first compared the peptidase activities of the asymmetric 20S CP with that of the symmetric 20S CP. We incubated separately asymmetric and symmetric 20S CP with a 10-fold stoichiometric excess of PA26PANc so that all 20S CP are either singly capped as PA26PANc-α_WT_ββα_K66A_ or doubly capped as PA26PANc-α_WT_ββα_WT_-PA26PANc, as confirmed by negative stain EM (Supplementary Figure 2A and B). Since PA26PANc opens the gate of the α-ring to which it binds, the doubly capped 20S CP has gates of both α-rings open, and the singly capped 20S CP has presumably one open gate in the α-ring that bears the activator (named proximal gate). Using LFP peptide as the substrate, the peptidase activity of singly and doubly capped proteasomal complexes were very similar (Figure 2A and B).

We further compared the ATP-dependent protein substrate degradation of GFP-ssrA by the asymmetric complex PAN-α_WT_ββα_K66A_ or symmetric PAN-α_WT_ββα_WT_-PAN. As with PA26PANc, we used excess PAN to ensure that all 20S CP are capped with activator (Supplementary Figure 2C and D). Because free PAN can unfold but not degrade GFP-ssrA, the time-dependent degradation of GFP-ssrA was measured not by decrease of GFP florescence, but instead by SDS PAGE gel and western blot analysis against GFP (Figure 2C and D). We found that PAN-α_WT_ββα_K66A_ and PAN-α_WT_ββα_WT_-PAN degrade GFP-ssrA at a similar rate. This result suggests that only one ATPase is needed to fully activate the 20S CP to unfold and degrade a folded protein substrate, or that two GFP-ssrA substrates are not unfolded simultaneously by the two PAN ATPases at the opposite ends of 20S CP.

One possible explanation of both the LFP peptidase activity and GFP-ssrA degradation results is that the distal gate remains closed in the asymmetric complex. In this view, having a single open gate in the 20S CP is not a rate limiting step for diffusive substrate entry (LFP, PA26PANc) or for active entry (GFP, PAN) and for proteolytic products to leave the degradation chamber. If so, the overall rate of peptide hydrolysis would not be greatly influenced by whether one versus two gates are open. An alternative explanation is that binding of an activator to the proximal α-ring is sufficient to allosterically open the gate of the opposite α-ring (named distal α-ring or gate). In this scenario, binding a single RP opens two gates. This idea was tested by determining the gate conformation of distal α-rings in asymmetric proteasome complexes.

### Gate conformation of distal α-ring in PA26PANc-associated asymmetric proteasomes

We determined a number of single-particle cryo-EM structures of the *in vitro* assembled asymmetric 20S CP with one activator bound, PA26PANc-α_WT_ββα_K66A_ and PAN-α_WT_ββα_K66A_. We did so in the presence or absence of substrates to test whether substrate processing would affect the outcome. When determining these reconstructions, we applied a C7 symmetry. The proximal and distal α-rings were identified easily using the bound activator as a reference. The resolution of all reconstructions was around 3Å, sufficient to reveal the gate conformations of each of the two α-rings, proximal and distal.

In eukaryotic 20S proteasomes, the α-ring contains seven homologous but different α-subunits, each with a different N-terminus. Through a complex hydrogen bonding network, these N-termini form a very ordered gate ^4^. In archaea 20S proteasome with homo-heptameric α- and β-rings, N-termini of the seven identical α-subunits also form a gate that prevents bulky substrates, such as long peptides or folded proteins, from entering the 20S proteasome for degradation ^18^. The N-termini of the seven α-subunits cannot form a structurally well-ordered gate through hydrogen bonds. In all structures of archaeal 20S proteasomes with a closed gate, the first nine residues of α-subunits are typically not resolved ^19,21^. NMR spectroscopy analysis of gate conformation of archaeal 20S shows that N-terminal tails of α-subunit point downwards in a closed conformation, and upwards in an open conformation ^24^. In our structural studies, we used the same criterion, tracing the pointing direction the N-terminal tails of the α-subunits to evaluate the gate conformation. This is based on previous single particle cryo-EM studies of gate opening in archaeal 20S proteasome and the NMR studies of archaeal 20S proteasome with open and closed gate (Supplementary Figure 3). This criterion is also used in a recent study of archaeal proteasomal complexes ^16^.

**Figure 3.**
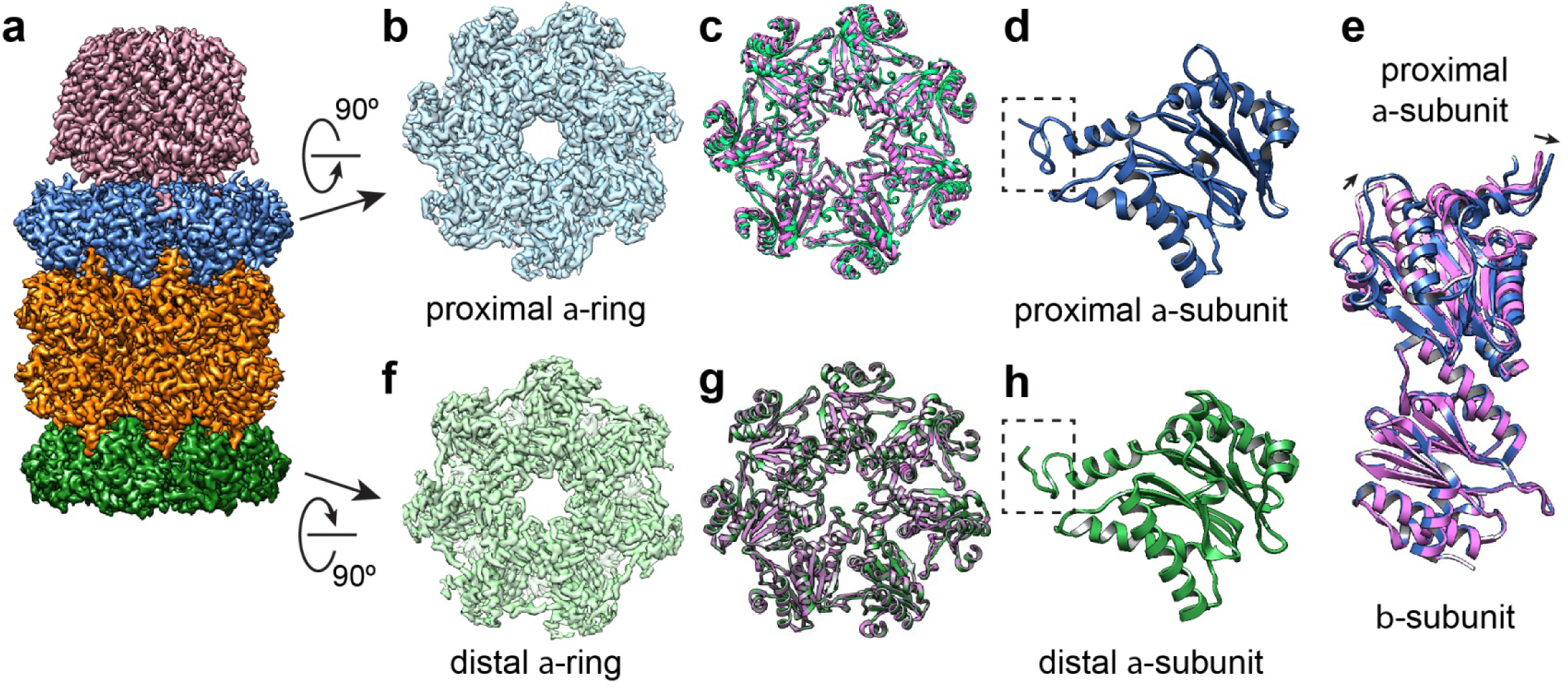
*Single particle cryo-EM structure of* PA26PANc-α_WT_ββα_K66A_ *complex*. **A**: Cryo-EM reconstruction of PA26PANc-α_WT_ββα_K66A_ at a resolution of 2.9 Å. Samples were prepared without substrate LFP peptide substrate present. **B**: Proximal α-ring of the 20S CP. **C**: Ribbon diagram of proximal α-ring (green) overlayed with that of the α-ring in closed conformation without bound activator (purple). Note that the two do not overlap, indicating rotation induced by activator binding. **D**: Ribbon diagram of α-subunits from proximal α-rings. The N-terminus of a-subunit within the dashed box points upwards in an open gate conformation. **E**: Ribbon diagrams of an α-subunit from the proximal α-ring and its interacting β-subunit (green) overlaid with the ribbon diagram of α/β-subunits from a 20S CP without activator bound (purple). Note that the β-subunit is perfectly aligned between the two, but α-subunit is rotated. **F**: The distal α-ring of the 20S CP. **G**: Ribbon diagram of distal α-ring (green) overlayed with that of the α-ring in closed conformation without bound activator (purple). Note that the two overlap nicely. **H**: Ribbon diagram of α-subunits from distal α-rings, where the N-terminus of a-subunit within the dashed box points upwards in an open gate conformation.

We first determined the structure of PA26PANc-α_WT_ββα_K66A_ to resolutions of 2.9 Å without LFP substrate and 3.4 Å with LFP (Figure 3, Supplementary Figure 4). In both structures, the conformation of the proximal α-ring and two β-rings match very well with the crystal structure of wild type symmetric 20S proteasome in complex with the same activator (PDB: 3IPM). As shown in this crystal structure ^23^, binding the hybrid activator PA26PANc to the α-ring causes α-subunit rotation, which opens the gate (Figure 3C-E). In both our structures (with and without LFP), the distal α-ring, that without an activator, also shows features that match an open gate, with all N-termini moved away from the pore and pointing upwards (Figure 3F and H). However, unlike the proximal gate, there is no rotation of distal α-subunits (Figure 3G). Rather, the conformation of this α-ring matches well with that in the crystal structure of archaeal 20S proteasome in complex with PA26 (PDB: 1YAR, ^21^), except that the gate is in an open conformation. The surprising observation that binding of PA26PANc to the proximal α-ring opens the gate of the distal α-ring is consistent with allosteric action. That the opening of the distal gate is independent of the presence of substrate implies that substrate presence or its active proteolysis within the β-ring does not mediate the observed allosteric effect.

**Figure 4.**
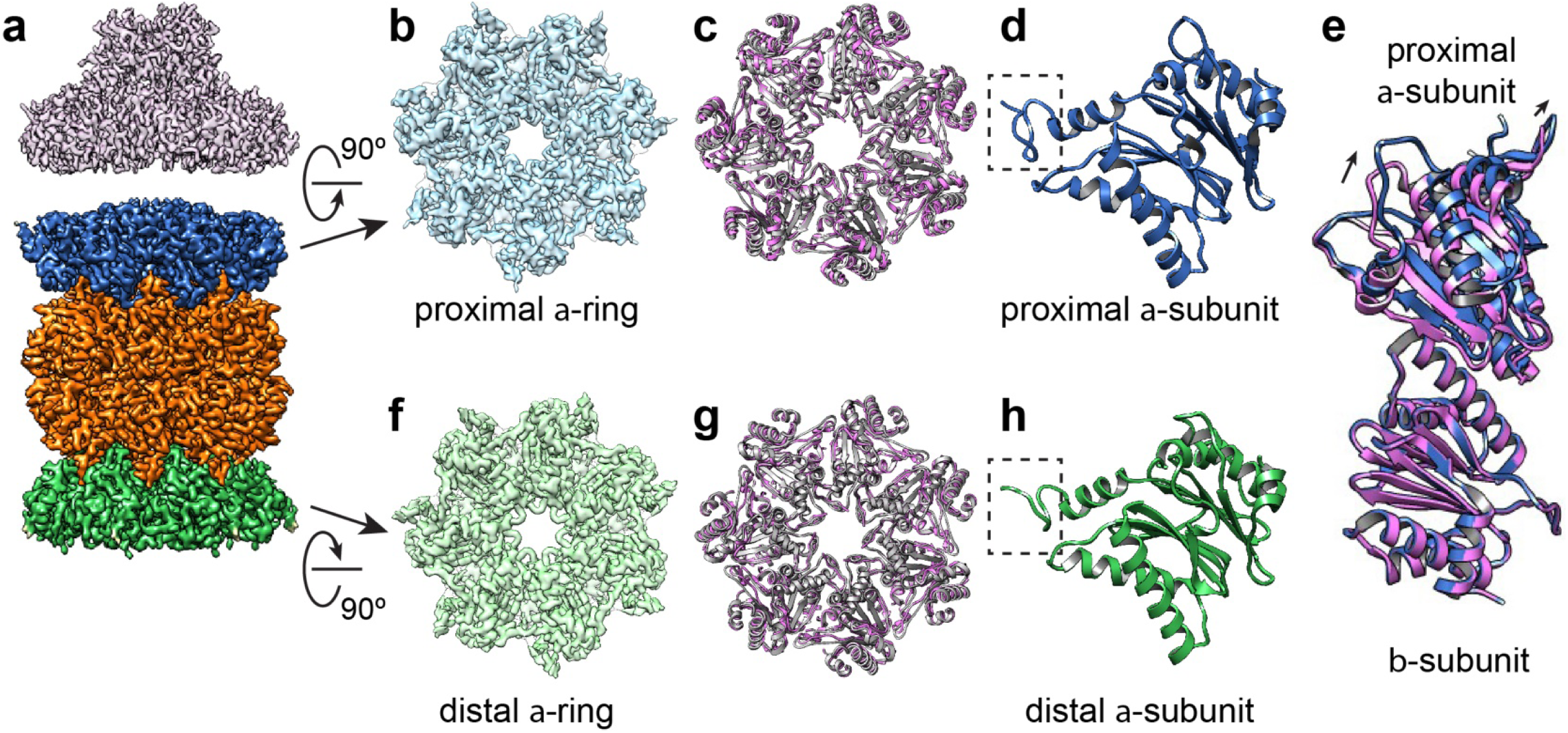
*Single particle cryo-EM structure of* PAN-α_WT_ββα_K66A_ *complex with GFP-ssrA substrate.* **A**: Cryo-EM reconstruction of PAN-α_WT_ββα_K66A_ with substrate GFP-ssrA at a resolution of 3.4 Å. **B** through **E** as in figure 3. **B**: Proximal α-ring of the 20S CP. **C**: Ribbon diagram of proximal α-ring (grey) overlayed with that of the α-ring in closed conformation without bound activator (purple). Note that the two do not overlap. **D**: Ribbon diagram of α-subunits from proximal α-rings. The N-terminus of α-subunit within the dashed box points upwards in an open gate conformation. **E**: Ribbon diagram of an α-subunit from the proximal α-ring and its interacting β-subunit (grey) overlayed with the ribbon diagram of α/β-subunits from a 20S CP without activator bound (purple). Note that the β-subunit is perfectly aligned between the two, and α-subunit is rotated. **F**: Distal α-ring of the 20S CP. **G**: Ribbon diagram of distal α-ring (grey) overlayed with that of the α-ring in closed conformation without bound activator (purple). Note that the two overlap nicely. **H**: Ribbon diagram of α-subunits from distal α-rings. The N-terminus of a-subunit within the dashed box points upwards in an open gate conformation.

### Gate conformation of distal α-ring in PAN-associated asymmetric proteasomes

We also determined structures of PAN-α_WT_ββα_K66A_ in the presence of ATP, without and with substrate GFP-ssrA, both to resolutions of 3.4 Å (Figure 4, Supplementary Figure 5). Because of the symmetry mismatch between the heptameric α-ring and hexameric PAN ATPase, PAN does not engage with α-ring symmetrically, and not every binding pocket in the CP is occupied by a C-terminus of PAN. Furthermore, engagement of PAN and α-ring is heterogeneous among different particles ^17^ and such heterogeneity limits the achievable resolution. To obtain sufficient resolution to evaluate the gate conformation, we applied a C7 symmetry. Thus, the density of PAN in this reconstruction does not represent its correct structure. Since it is not the emphasis of this study, we did not attempt to obtain a high-resolution structure of PAN bound with the 20S CP. The conformation of proximal gate (and presumably the distal gate as well) is an average of slightly different conformations of α-subunits within the ring. Even so, the conformation of the gate in the C7 symmetrized density map still represents the state of the gate. Whether without or with the substrate, the proximal gates are in an open conformation, consistent with the gate opening by the C-termini of PAN with its C-terminal HbYX motif. Furthermore, the C7 symmetry averaged density map still shows rotation of α-subunits induced by binding of PAN’s C-termini. This result (Figure 3) using PAN is consistent with the structures determined with PA26PANc. Thus, structures of PA26PANc-α_WT_ββα_K66A_ and PAN-α_WT_ββα_K66A_ both show that the gate of the distal α-ring is open even without direct association with an activator; this distal gate is opened allosterically by the engagement of an activator to the proximal α-ring. These data indicate that an RP carrying the HbYX motif, which intercalates within the pockets between neighboring pairs of α-units, and induces a small rotation in the α-unit to open the gate, can mediate allosteric gate opening.

**Figure 5.**
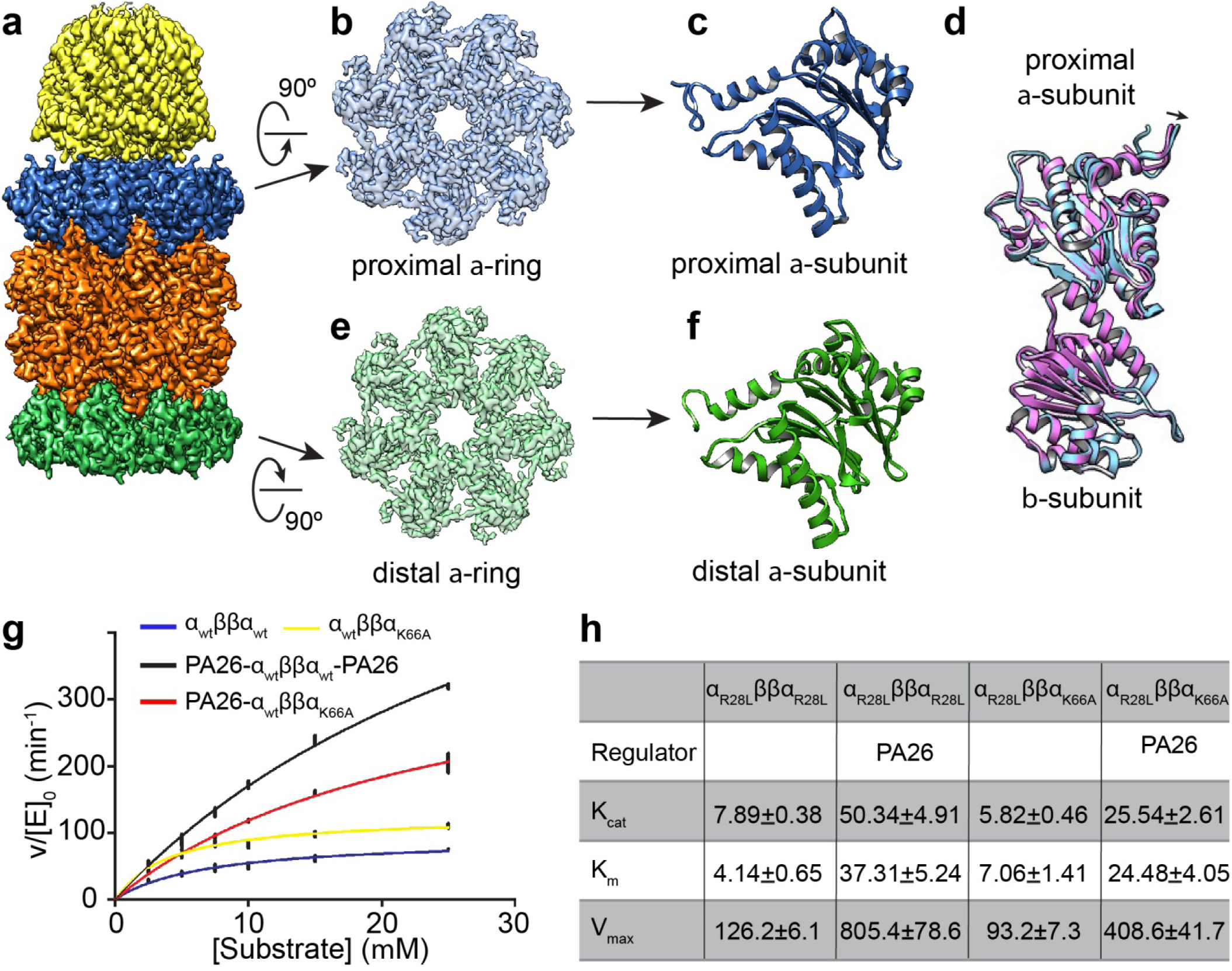
*Single particle cryo-EM structure of* PA26-α_WT_ββα_K66A_ *complex without substrate.* **A**: Cryo-EM reconstruction of PA26-α_WT_ββα_K66A_ at a resolution of 3.5 Å. **B and E**: Proximal and distal α-ring of the 20S CP. **C** and **E**: Ribbon diagram of α-subunits from proximal and distal α-rings. Note that the N-terminus of proximal α-subunit points upwards, consistent with an open gate conformation. In contrast, the N-terminus of distal α-subunit points downwards, consistent with a closed gate conformation. **D**: Ribbon diagram of an α-subunit from the proximal α-ring and its interacting β-subunit (grey) overlayed with the ribbon diagram of α/β-subunits from a 20S CP without activator bound (purple). Note that both α- and β-subunits are well-aligned between the two. **G**: Kinetics of LFP degradation by symmetric (blue) and asymmetric (yellow) 20S CP alone and with double (black) and single (red) activator PA26. **H**: Table summarizes the *k*_*cat*_, *K*_*M*_, and *V*_max_ kinetic parameters of peptidase activity of symmetric and asymmetric 20S CP without and with PA26 activator.

### Gate of distal α-ring in asymmetric proteasomes using PA26 without HBYX motif

To test whether the HbYX motif is a required element for inducing allosteric communication between the two distant α-rings, we further determined the structure of the asymmetric CPα_WT_ββα_K66A_ in complex with a single PA26, which does not have an HbYX motif in its C-terminus. As expected, the gate in the proximal α-ring is open and there is no rotation of the α-subunit, but the activation loop of PA26 is engaged with the reverse turn loop of 20S CP (Figure 5, Supplementary Figure 6). In contrast with the results using PAN and PA26PANc, in an asymmetric proteasome bearing PA26, the gate of the distal α-ring is closed. This result suggests that binding a proteasomal activator without a C-terminal HbYX motif to one α-ring opens the proximal gate only, but does not induce allosteric opening of the distal gate.

These results motivated us to further test the peptidase activities of asymmetric 20S CP with a single capped PA26, in comparison with that of the symmetric 20S CP with two PA26 caps. If the diffusion rate of LFP peptide through the open gate into 20S CP is a rate limiting step, the peptidase activities of symmetric 20S CP double capped with PA26 (two gates open) should be higher than that of the asymmetric 20S CP capped with only one PA26 (one gate open). Indeed, the peptidase activity of symmetric 20S CP doubly capped by PA26 is about twice that of the singly capped asymmetric CP (Figure 5B), as predicted from the structures and the stated hypothesis. This result further suggests that the rate of unfolded peptide substrates diffusing through an open gate into the proteolytic chamber is a rate limiting step for the peptidase activity of the 20S proteasome.

### Cooperative binding associated with distal gate opening

Binding of a regulatory cap to an α ring promotes local gate opening. In case of activator with a C-terminal HbYX motif, such as PAN or PA26PANc, the distal gate is also opened allosterically. Such an unoccupied α ring with an allosterically opened gate, in which N-termini of all α subunits are moved away from the pore and pointing upwards, may kinetically favor activator binding over an α ring that is otherwise identical but with its gate closed. The empty and open distal gate is preconfigured to accept an activator. This hypothesis makes a testable prediction: PAN or PA26PANc binding to 20S CP is cooperative, but PA26 binding to 20S CP is not cooperative. Cooperative binding of an RP implies that the distribution among uncapped, singly capped and doubly capped particles will show a deficit of singly capped 20S CP, and a corresponding increase in uncapped and doubly capped particles. Interestingly, such cooperative binding of eukaryotic 19S RP to 20S CP has been described for the eukaryotic proteasome ^25^.

To test this hypothesis, we collected electron micrographs of negatively stained wild type symmetric 20S CP incubated with either PA26PANc or PA26 in two different molar ratios, 4:1 and 3:1. We picked all 20S CP particles, with or without caps, and classified them into three categories: 20S CP alone and with one or two activators bound (Supplementary Figure 7). The probability of a cap binding to an α ring, P, was estimated from the number of particles in each category. The hypothesis of non-cooperative binding of cap to 20S CP predicts a binomial distribution of particles in each category: P^2^ for doubly bound 20S CP, 2P(1-P) for singly bound 20S CP, and (1-P)^2^ for unbound CP (Supplementary Table 3 and 4). Comparing the observed and the predicted number of singly capped particles, singly capped PA26PANc is significantly less than that predicted by the null hypothesis. Such a deficit of singly capped complexes is the hallmark of positive cooperativity. The null hypothesis was rejected with high probability by χ^2^ analysis. However, a similar analysis with PA26 revealed no evidence of positive cooperativity. The numbers of observed particle in all three categories are quite close to the prediction, supporting the null hypothesis of no cooperativity. Overall, these results support the conclusion that a preformed unoccupied open gate in the α ring has increased affinity for binding RP. Cooperative RP binding and distal gate opening are thus seen to be different aspects of a single underlying mechanism.

## Discussion

In both eukaryotes and archaea, the 20S CP is two-fold symmetric, with identical α-rings at each end; these can bind to identical or different proteasomal activators. In the case of the most widely studied eukaryotic 26S proteasome, two 19S activators are symmetrically attached to the 20S CP. However, a significant fraction of 20S proteasome particles of eukaryotic cells are capped with only one 19S activator, forming asymmetric proteasomal complexes. In addition, there are also 20S CP capped asymmetrically by a 19S RP ATPase activator and a non-ATPase activator, such as 11S PA28-hybrid proteasomes ^7^. It has been unclear whether proteasomes function similarly with one or two caps. This has been difficult to test biochemically, because homogeneous native proteasomal complexes with only one activator cannot be readily purified. Our *in vitro* assembled asymmetric 20S CP enabled us to generate homogenous singly capped proteasomal particles. The biochemical data presented here show that, when bound with activator that has an HbYX motif at its C-terminus, single and double capped 20S CP have similar substrate degradation rates, both for small peptide substrates and protein substrates that require ATPase-dependent unfolding. However, when bound with PA26, which does not have a C-terminal HbYX motif, peptidase activity of doubly capped proteasomes is greater than for those with a single cap. Consistently, our structural and cooperativity binding analyses revealed that allosteric opening of the distal gate by engagement of an activator to the proximal α-ring and cooperative binding of an activator to the 20S CP are observed when an HbYX motif is present in the C-terminus of the activator.

In archaea, only proteasomal ATPases (PAN or Cdc48) are known to contain such a native HbYX motif in its C-terminus ^26^. As shown here, that motif functions similarly when artificially grafted to the C-termini of the non-ATPase activator PA26. Previous studies showed that binding of an activator with a C-terminal HbYX motif causes a rotation of the α ring ^19^. The capacity of the HbYX motif to allosterically open the distal gate suggests that the rotation may play a role in initiating the allosteric action. The present observation, that engaging an activator to one α-ring is propagated to the opposite α-ring, is remarkable, as it implies action at a great distance, across the full span of the CP, some 150Å. The allostery must necessarily be transmitted by a series of conformational changes within the β-rings separating the non-contiguous α rings in the CP. However, except for the changes of N-terminal gating residues of α-subunits, we failed to detect any noticeable conformational change in the β-rings sandwiched between the two α-rings. Therefore, how the β-rings transmit the conformational changes from the proximal α-ring to the distal one remains to be determined.

Previous descriptions of allosteric interactions within the eukaryotic proteasome CP predominantly focused on how the conformational changes associated with substrate engagement with β ring is propagated to the α ring and alters its interaction with RP ^27-31^. The findings described here extend that allosteric coupling across the full expanse of the CP, from one α ring to the other at the opposite end. Distal gate opening is observed regardless of whether substrate is present and being processed.

Allosteric communication between two oppositely positioned α rings is reflected by two sorts of observation: cooperative binding and distal gate opening by an ATPase regulatory particle with an HbYX motif. These forms of regulation can be understood mechanistically as one: A preformed open gate in an α-ring confers a higher affinity to an ATPase RP than does an α-ring with a closed gate, thus promoting the formation of a proteasomal complex with doubly capped ATPase. The two findings-cooperativity and distal gate opening-may thus be regarded as related manifestations of a common underlying mechanism.

We propose that the mechanism explanatory of our observations of PA26PANc cooperative binding suggests that a non-ATPase RP, one without a C-terminal HbYX motif, will also have a higher affinity to an α-ring with a preformed open gate. If so, mixing 20S CP with a combination of ATPase and non-ATPase activators would bias the population of proteasomal complexes towards either those doubly capped with ATPases, or hybrid complexes with an ATPase and a non-ATPase at the two opposite ends. Within eukaryotic cells, such cooperativity might promote for example the formation of either 26S doubly capped with two 19S RP or hybrid proteasomes capped with a 19S RP and a non-ATPase activator, such as 11S or PA200, and at the same time minimize formation of proteasomes doubly capped with two non-ATPase activators. Such proteasomal particles with doubly capped non-ATPase activators have no obvious functionality in degrading protein substrates.

From the prospective stated, RPs may be regarded as falling into two classes-those that can act as both transmitters and recipients of allosteric information (as here documented for RPs with an HbYX motif) and those that are solely recipients (RPs that lack the motif). Proteasomes are modular molecular machines. The presence within an individual cell of multiple alternate RPs, some functioning dually as transmitters and recipients of allostery, others solely as recipients, enhances the potential number of stable CP-resident molecular species, thus regulating how CP pools partition among these alternate populations. Concentrations of the individual RP modules and their absolute and relative affinities for the CP α ring must be important determinants of how a CP population is partitioned among alternate forms. Cooperative binding provides an additional form of regulatory complexity.

We have examined cooperativity in a highly constrained experimentally accessible system, looking at interactions among pure populations of archaea 20S in the presence of a single species of CP, here specifically PA26PANc or PA26. In this simple context, cooperativity has a simple consequence: a deficit of singly capped species and concomitant excess of uncapped and doubly capped. Our assays of various biochemical activities have revealed that adding a second identical cap provides little enhancement of activity, at best two-fold. Such a small change in biochemical outcome seems an implausible foundation to support a conserved complex allosteric interaction, one common to both eukaryotes and archaea. Cooperativity more plausibly promotes formation of proteasome species with hybrid capping, in which distinct RP complexes inhabit the same proteasome complex, for both archaea and eukaryotes. Hybrid asymmetric proteasomes are abundant in cells, but have unclear function. The mechanism of their formation and maintenance is unexplored. Allosteric α ring coupling may provide a means for fine tuning hybrid formation.

Certain archaea species lack an obvious homolog of PAN. It has been conjectured that they utilize an alternate ATPase activator, homologous to the Cdc48/p97 ATPase of eukaryotes^26^. Cdc48/p97 contains the C-terminal HbYX motif. It would be of interest to determine whether that activator can also confer distal allosteric gate opening.

Our studies were carried out using archaea 20S proteasome, because it is feasible to assemble asymmetric 20S CP *in vitro*. Nevertheless, our findings could also shed light on the eukaryotic proteasomal complex, as previous biochemical and structural studies demonstrate similarities between the two. Certainly, eukaryotic proteasome are more complex. Only three of six Rpt subunits contain the HbYX motif, and full gate opening of the proximal α-ring depends on engagement of substrates ^32^. At present, current single particle cryo-EM structural studies of asymmetric eukaryotic proteasomes have not determined whether the distal gate or distal α-ring is allosterically influenced by the functional state of the proximal activator, as the full array of experimental conditions relevant to this question have not been explored. Further investigation would be required to fully address the questions posed here.

## Supporting information

Manuscript BioRxiv.pdf

## Acknowledgement

We thank H.M. Kim and K. Egami for their efforts in the early stage of this project, and M. Braunfeld at UCSF cryo-EM facility for technical support. This work was supported in part by grants from National Institute of Health (R01GM082893, S10OD021741 and S10OD020054 to Y.C. and R01GM107124 to P.C.). Y.C. is an Investigator with the Howard Hughes Medical Institute.

## Author contributions

Z.Y. performed most of the biochemical and cryo-EM experiments and participated in experimental planning. Z.Y and Y.Y. designed mutagenesis experiments. F.W. and Z.Y. developed a modified EM grid to avoid preferred particle orientations. Z.Y. and A.G.M. collected initial high-resolution datasets. P.C. and Y.C. planned experiments, supervised experiments and performed data analysis. Z.Y., P.C. and Y.C. contributed to manuscript preparation.

## Data availability

All cryo-EM density maps and corresponding coordinates are deposited and available in databases with access numbers: EMDB-20881 and 6UTJ (singly capped PA26PANc-20S complex), EMDB-20879 and 6UTH (singly capped PA26PANc-20S complex with substrate LFP), EMDB-20878 and 6UTG (singly capped PA26-20S complex), EMDB-20880 and 6UTI (singly capped PAN-20S complex), EMDB-20877 and 6UTF (singly capped PAN-20S with substrate GFPssrA).

## Methods

### Protein expression and purification

Recombinant wild type and mutant forms of *Thermoplasma acidophilium* 20S core particles (CP), its individual α- and β-subunits, and various proteasomal activators (PAN ATPase from *Methanococcus jannaschii*, and PA26 from *Trypanosoma brucei*) with various affinity tags were expressed in *E. coli* BL21(DE3), using plasmids pET28a, pET15b or petDuet-1 (Supplementary Table 1). A TEV proteolytic site was inserted adjacent to tags for tag removal after final purification and complex assembly. After culture in LB broth at 37°C, expression was induced with 0.5mM IPTG at a cell density of OD 0.6-1.0, and cells further cultured at 18°C overnight before harvest. Cells were lysed by emusiflex, microfuidizer or sonication and affinity purification performed using methods appropriate to the specific tags. Purification of individual proteins was assessed by SDS PAGE and Coomassie staining. Fully functional symmetric 20S CP was assembled by mixing individually expressed α- and β-subunits. Residues 2-8 of the β-subunit were truncated to simulate the autoproteolytic N-terminal pro-peptide truncation that occurs during native CP assembly.

### In vitro reconstitution of asymmetric 20S CP and its complex with activators

*In vitro* assembly and purification of 20S CP with the intended compositions were performed using a multi-step affinity purification. Specifically, two different types of individually purified α-subunits were incubated with β-subunit in a 1(α type 1):2(β):1(α type 2) molar ratio in Tris buffer at pH7.5 for 1 hour at 37°C. Typically, the mixture contains different unassembled single or double α-rings (α_WT_, α_K66A_, α_WT_α_WT,_ α_K66A_α_K66A,_ α_wt_α_K66A_), symmetric, asymmetric 20S CP and free β monomers. Affinity purification with streptrap column selects all assembled or individual α-subunit containing α_K66A_. After streptrap column, a TEV protease enzyme was used to cleave all the tags except MBP, followed by another affinity purification by MBP column, which yields fully assembled asymmetric 20S CP. The functionality of assembled 20S CP was confirmed by peptidase activity.

To assemble complexes of asymmetric 20S CP with PA26PANc or PA26V230F, regulatory particles were added to partially purified pre-assembled asymmetric 20S CP before affinity purification by MBP column in stoichiometric excess (10:1, RP:CP). An MBP column was used to capture fully assembled asymmetric 20S CP in complex with RP. To assemble the complex of 20S CP with PAN, in vitro assembled 20S CP, either symmetric or asymmetric, was mixed with PAN in stoichiometric excess (4:1, PAN:CP) in the presence of nucleotide. For ATP dependent GFP-ssrA degradation assay, ATP was used. For structural study of the 20S-PAN complex, ATPγS was used. Final complex products were checked by negative stain EM to assess whether they had the anticipated structure.

### Peptidase activity assay

Peptidase activity (Figure 2A and 5G) was determined by measuring the fluorescent intensity generated by LFP hydrolysis, a fluorogenic 9-residue peptide, as described previously ^18^. Assays were performed in 200 μl buffer (25 mM Tris pH 7.5, 10 mM MgCl_2_, 5% glycerol). In a typical assay, 1pmol 20S proteasome and 7.14pmol PA26 or its derivative proteins RP (in a final molar ratio of approximately 7:1) were incubated with LFP in various concentrations, ranging from 2.5 μM to 25μM LFP. Reactions components were assembled on ice and assays initiated by heating to 37°C. Fluorescence emission data (322/398 nm) were collected on a plate reader (SpectraMax M5). For each condition three independent measurements were performed. For each reaction, after an initial period of thermal equilibration, degradation rates were determined by least squares fitting of a linear equation to the time dependent data ^18,23^.

### GFP-ssrA degradation

An eleven residue ssrA peptide, AANDENYALAA, appended to the C-terminus of GFP provides a tag recognized by PAN for degradation ^33^. 50nM 20S proteasome and 200nM PAN were mixed in the presence of 5mM ATP, in reaction buffer of 25mM Tris pH7.5, 150mM NaCl, 10mM MgCl_2_ and 5% glycerol in a volume of 60μl. Reactions were initiated by addition of GFP-ssRA to a final concentration of 1μM. 10μl aliquots were collected at specific time points and immediately mixed with SDS loading buffer and heated to 95 degrees to terminate enzymatic degradation. To determine the rate and extent of GFP-ssrA degradation, samples were resolved by 4-15% SDS-PAGE for either Coomassie blue staining or western blot by anti-GFP (a kind gift from Xi Liu and John Gross) to visualize the specific GFP bands.

### Electron-microscopy data acquisition

For negative staining, 2.5 µl of the purified proteasome without or with its regulator (~0.01-0.1 mg/ml) was applied to glow-discharged EM grids covered by a thin layer of continuous carbon film (TED Pella, Inc.) and stained with 0.75% (w/v) uranyl formate solution as described ^34^. EM grids were imaged on a Tecnai T20 microscope (Thermo Fisher Scientific) operated at 200 kV with a TVIPS TemCaM F816 scintillator-based CMOS camera (TVIPS). Images were recorded at a magnification of 50,000x, which resulted in a 3.139 Å pixel size on the specimen. Defocus was set to −1.5 µm to −2 µm. For cryo-EM, 2.5 µl of purified proteasome sample (~1.0 mg/ml) was applied to glow-discharged holy carbon grids (Quantifoil 400 mesh Cu R1.2/1.3). For PAN-T20S sample, 2.5μl (~0.1 mg/ml) was applied onto amino-functionalized graphene oxide holy carbon grids (Quantifoil 300 mesh Au R1.2/1.3). The grids were blotted by Whatman No. 1 filter paper and plunge-frozen in liquid ethane using a Mark III Vitrobot (Thermo Fisher Scientific) with blotting times of 8-12 seconds at room temperature and over 90% humidity. Cryo-EM datasets were collected using SerialEM in three different microscopes all equipped with Field Emission source and K2 or K3 camera (Gatan Inc.) operated in super-resolution mode (Supplementary Table 2).

### Image processing and model building

For all negative stain EM datasets, RELION was used for particle picking and 2D classification. For all cryo-EM datasets, movies were motion-corrected by MotionCor2 ^35^. Motion-corrected sums without dose-weighting were used for defocus estimation by using GCTF ^36^. Motion-corrected sums with dose weighting were used for all other image processing. All datasets were processed similarly. In summary, particles were picked automatically by Gautomatch (http://www.mrclmb.cam.ac.uk/kzhang/Gautomatch/), extracted and 2D class averages by RELION and then imported into cryoSPARC for 3D classification and refinement procedures. Crystal structure (PDB: 3IPM) lowpass filtered to 30 Å was used as an initial model. Particles were sorted by iterative 2D classification and heterogenous refinement. Final maps were refined and reconstructed by 3-D auto-refinement. Resolution was estimated using FSC=0.143 criterion^37^. Atomic model of *Thermoplasma acidophilium* 20S proteasome (PDB: 3IPM) was used as a starting model followed by manually adjusting in COOT to fit the density map, followed by iterative refinement with Phenix.real_space_refine and manually adjustment in COOT. This process was repeated until Ramachandran validation was satisfied. Chimera ^38^ and Pymol ^39^ were used to prepare images.

### Cooperativity of regulatory complex binding

140nM of wild type *Thermoplasma acidophilium*as 20S CP was mixed with PA26PANc or PA26V230F at various molar ratios in a total volume of 10μl and incubated at 37°C for one hour in 25mM Tris buffer of pH 7.5 and 150mM NaCl. To prepare for negative stain EM grids, the mixture was diluted 3-fold to provide adequate particles distribution. Particle picking was performed by combining automatic (using Gautomatch) and manual picking to ensure all separated particles in each micrograph being analyzed are picked. Relion was used for 2D classifications to separate singly-, doubly-or non-capped proteasomes. The numbers in each class were tabulated for cooperativity calculations. Micrographs of each sample were randomly and equally divided into three groups to determine reproducibility of determinations of particle composition.

To assess the cooperativity of RP binding to 20S CP, the experimental distribution of particles among doubly, singly and uncapped complexes was compared to the distribution assuming non-cooperative binding, the null hypothesis. Based on the fraction of 20S CP particles with double (E2), single (E1) and no cap (E0), the probability of a α ring being capped by an activator is P = (2 × E2 + E1)/2, and being not capped is Q = (2 × E0 + E1) / 2 = 1 – P. Then, according to the null hypothesis, capping of each α ring in a 20S CP is independent or not cooperative, the anticipated numbers of particles with double, single and no cap are P^2^, 2PQ and Q^2^. Experimental and calculated distributions were compared using χ^2^ analysis to determine whether the experimental particle distribution was significantly different from that predicted by the null hypothesis. The results of all analyses are listed in Table 1 and Supplementary Table 3 and 4.

**Table 1:** Cooperative association of 20S CP with PA26PANc or PA26

| Sample: | Observed<br>2/1/0 caps<sup>a</sup> | Calculated<br>2/1/0 caps<sup>b</sup> | $\chi^2$<br>2 degrees of freedom | P |
| --- | --- | --- | --- | --- |
| PA26PANc:20S(4:1) | 4005/7767/8562 | 3070/9659/7625 | 383 | $<< 10^{-8}$ |
| PA26PANc:20S(3:1) | 1091/4623/16808 | 518/5788/16238 | 344 | $<< 10^{-8}$ |
| PA26:20S (4:1) | 6624/8591/2233 | 6840/8165/2443 | 23 | $< 10^{-5}$ |
| PA26:20S (3:1) | 2318/9720/7956 | 2439/9077/8477 | 42 | $< 3 \times 10^{-8}$ |

20S CP was incubated with PA26PANc or with PA26 at the indicated molar ratio of 4:1 or 3:1, and the number of particles with two, one or no RP caps was determined by negative stain EM. The number of each species theoretically expected, assuming association was not cooperative, was calculated, as described in Methods. The particle number distributions for the observed and calculated were used to calculate the $\chi^2$ statistic and its associated P value.
<sup>a</sup>Observed number of doubly capped, singly capped and uncapped particles.
<sup>b</sup>Calculated number of doubly capped, singly capped and uncapped particles, calculated using the binomial distribution, assuming binding at each end of 20S CP is independent.

