## Supplementary material for "Allosteric coupling between α-rings of the 20S proteasome": Manuscript BioRxiv.pdf

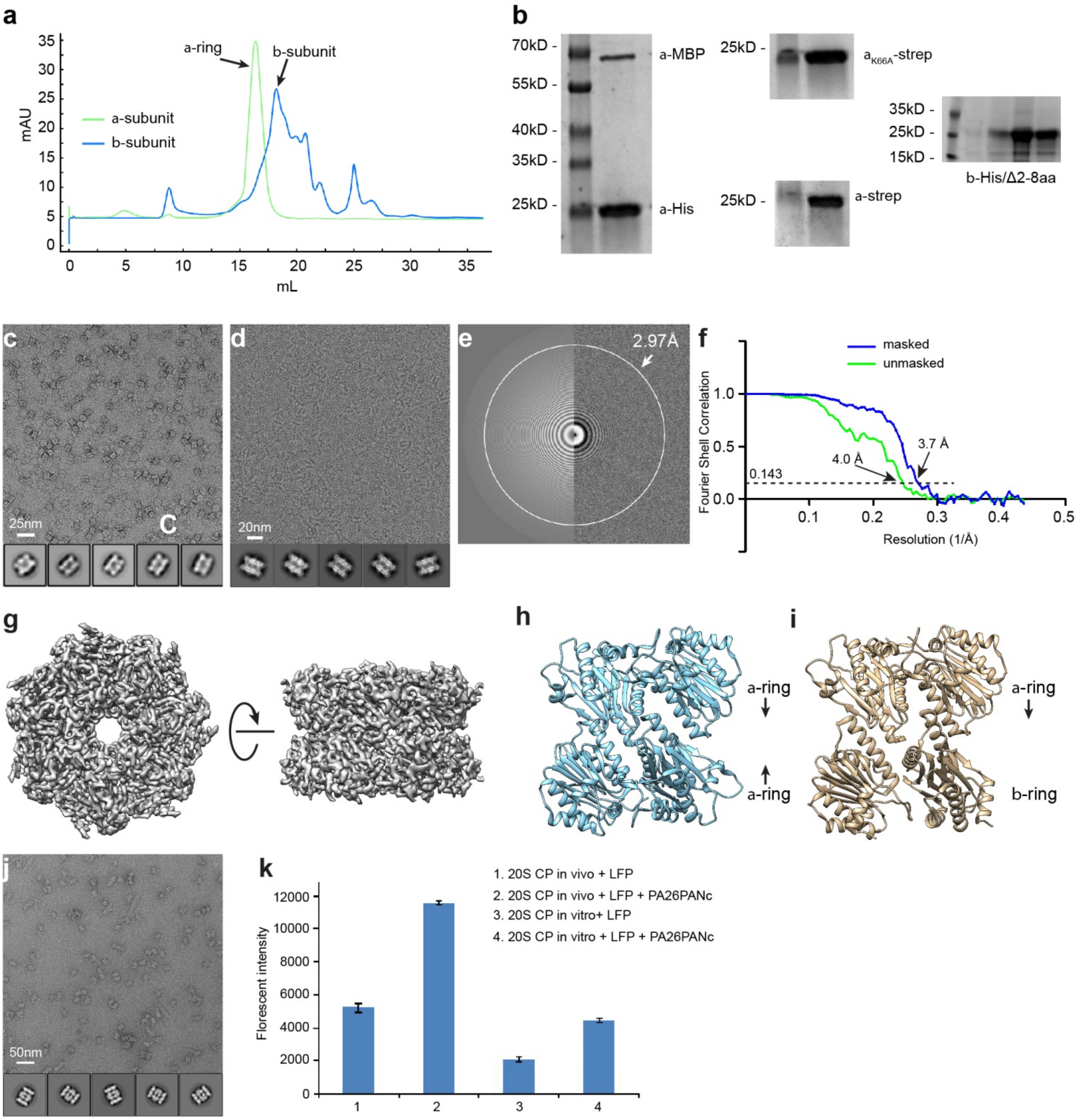


### Yu et al Supplementary Figure 1

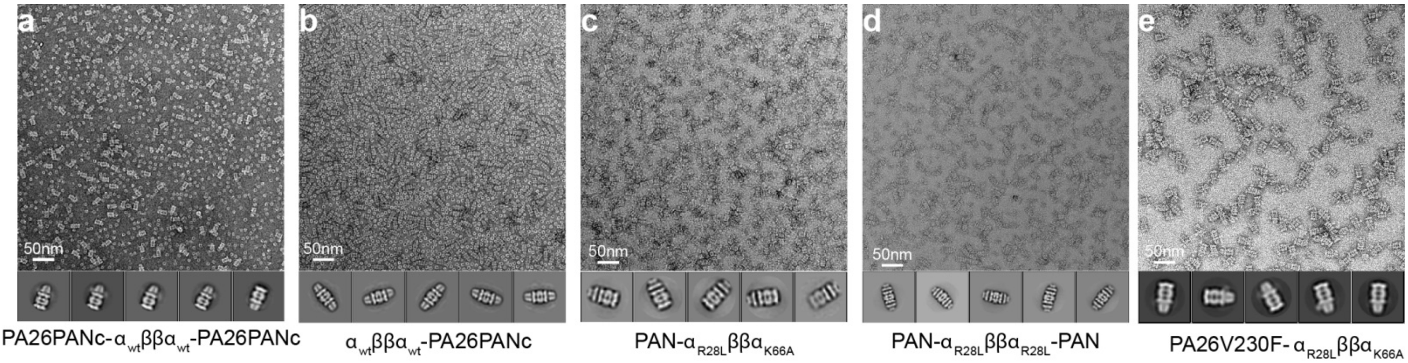


### Yu et al Supplementary Figure 2

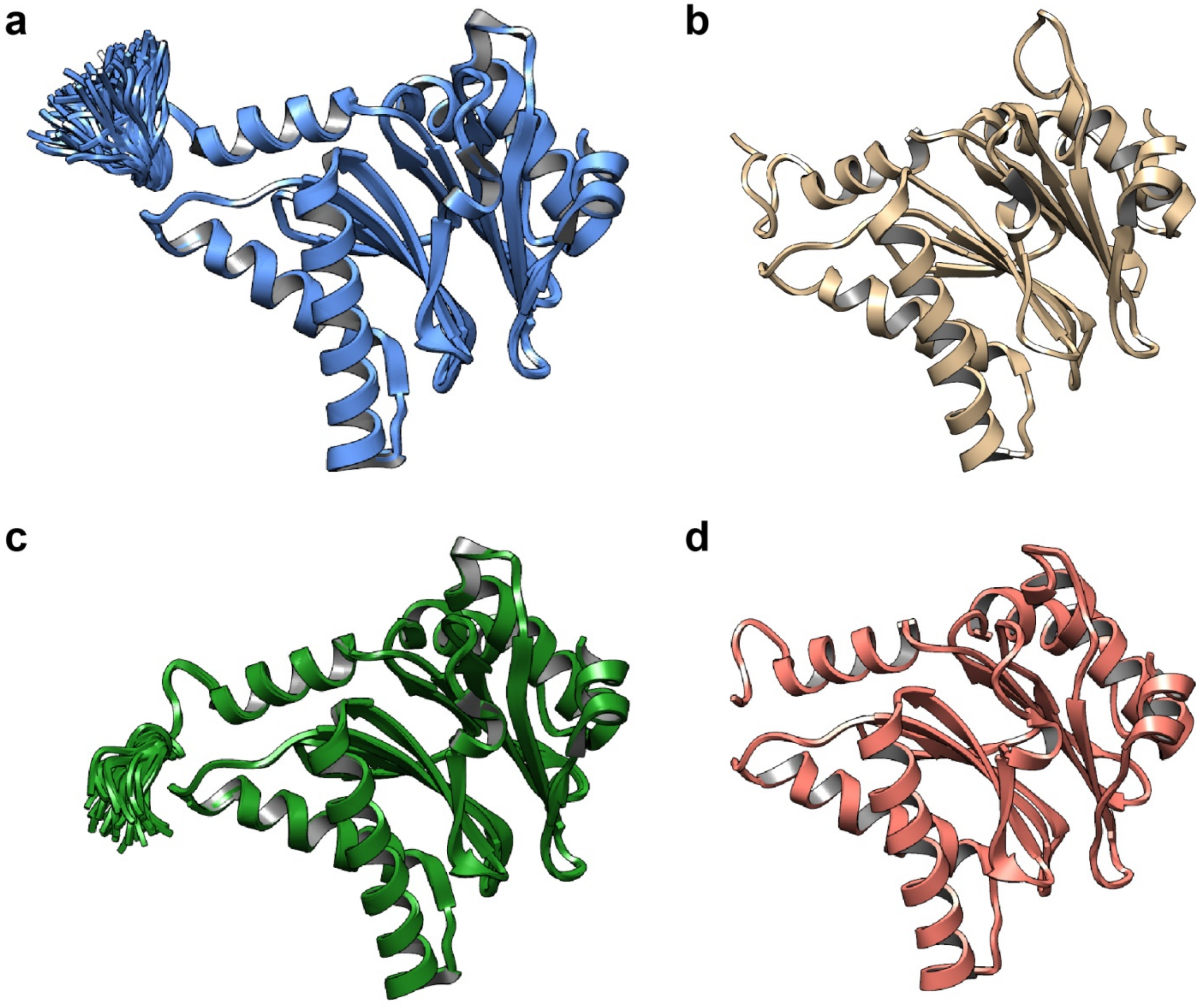


**Yu et al Supplementary Figure 3**

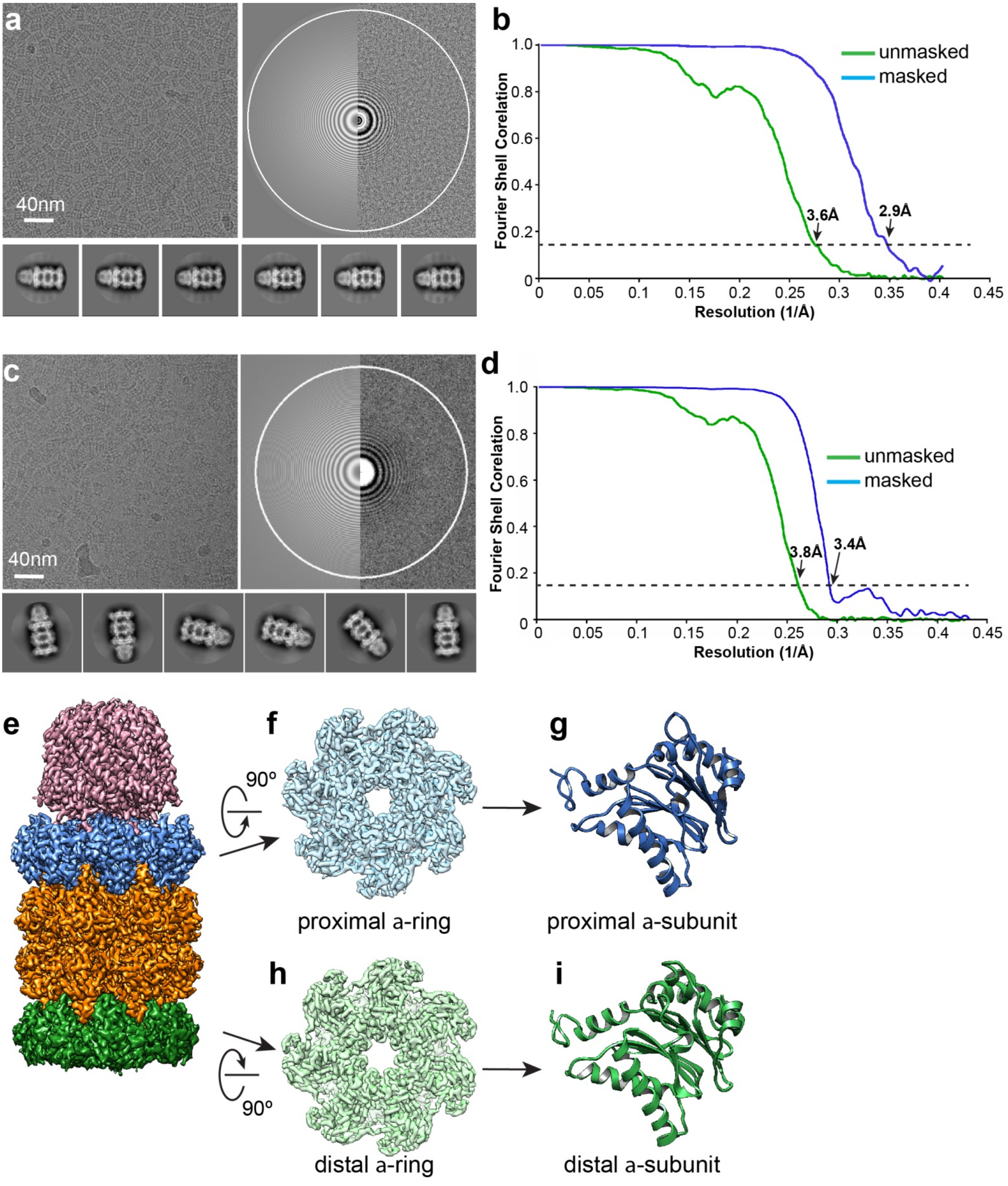


**Yu et al: Supplementary Figure 4**

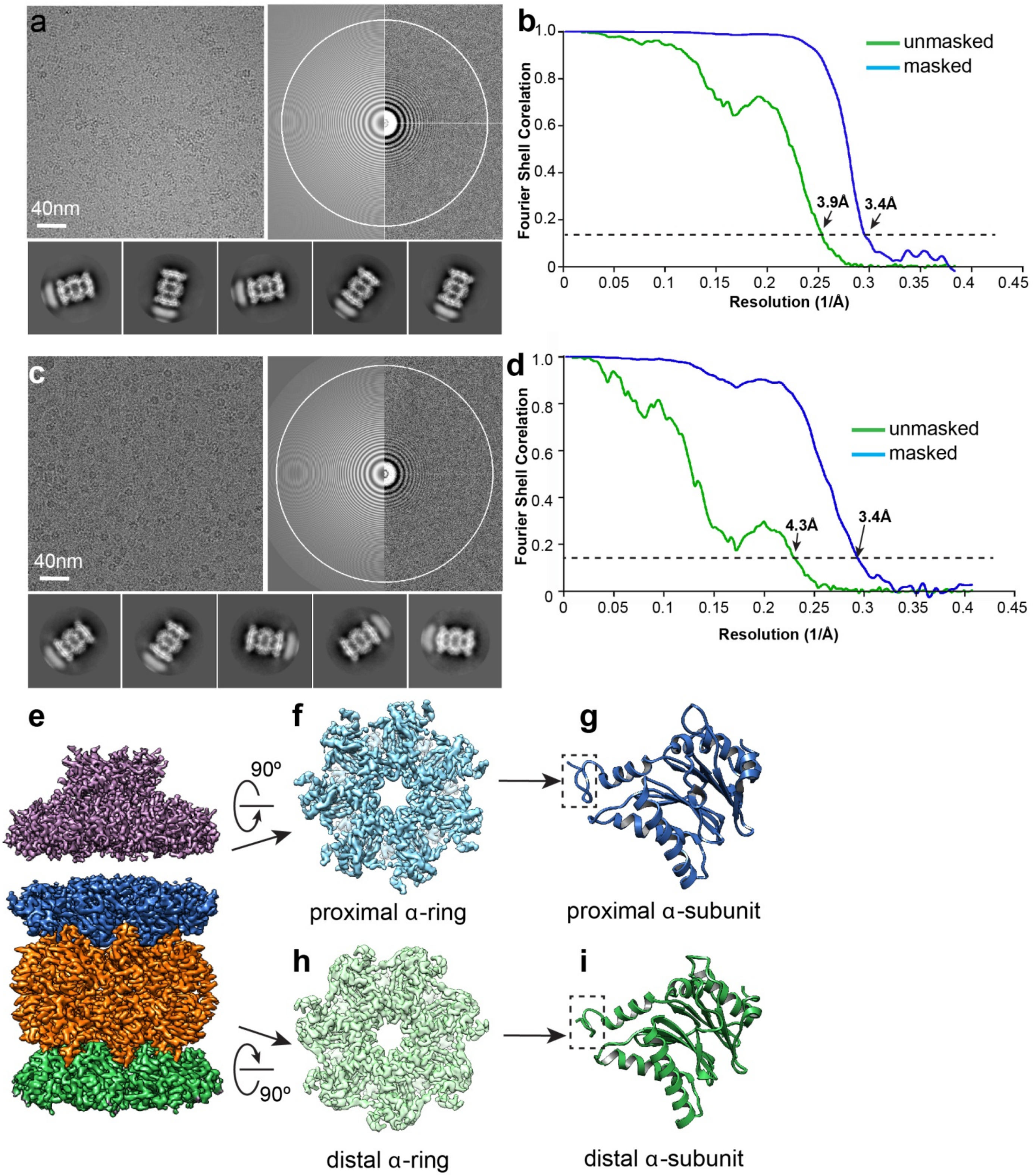


**Yu et al: Supplementary Figure 5**

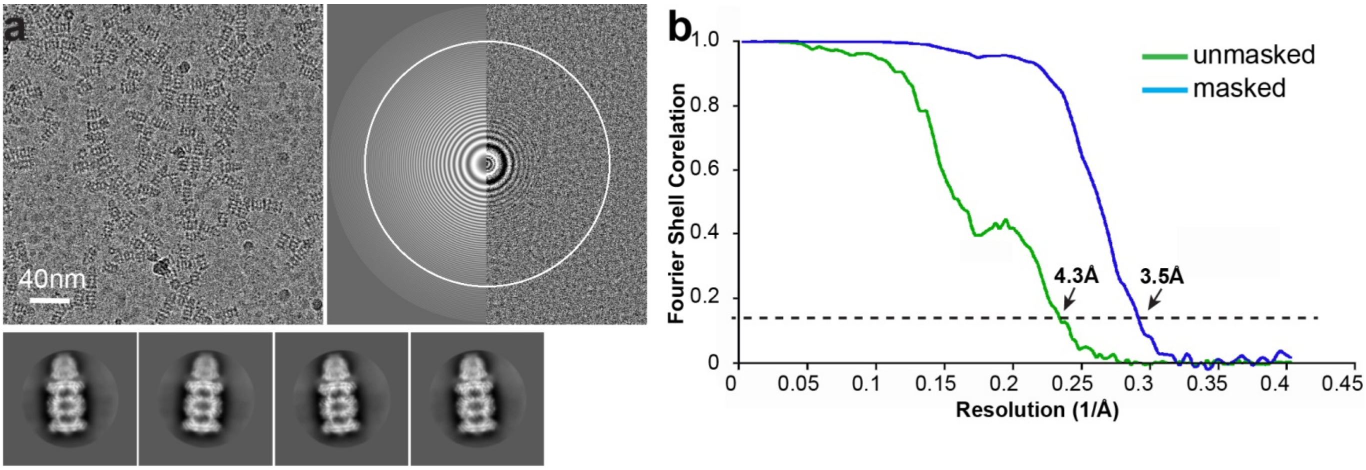


**Yu et al: Supplementary Figure 6**

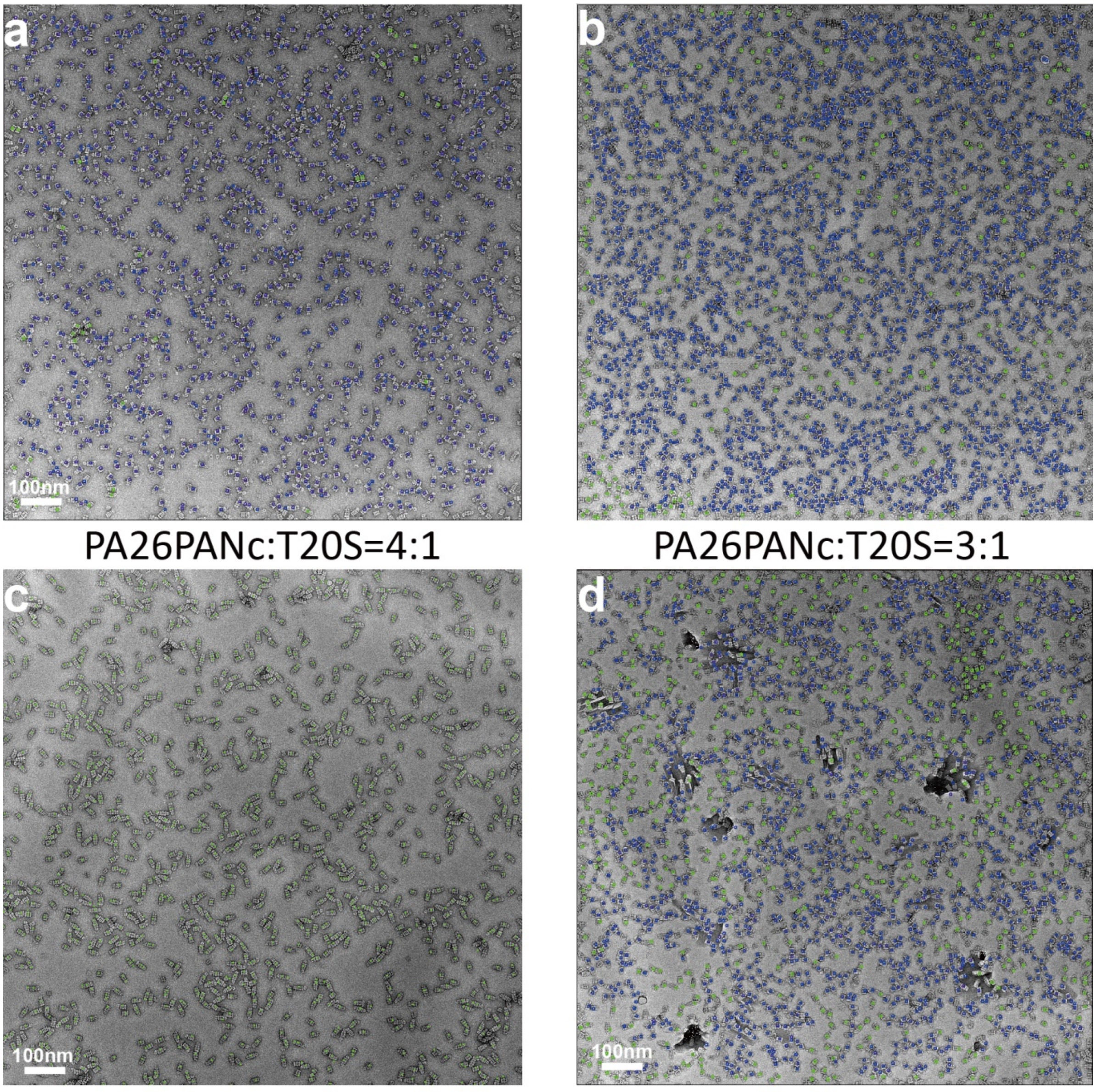


**Yu et al: Supplementary Figure 7**
